# CREB is Required in Excitatory Neurons in the Forebrain to Sustain Wakefulness

**DOI:** 10.1101/2020.09.22.308403

**Authors:** Mathieu E. Wimmer, Cui Rosa, Jennifer M. Blackwell, Ted Abel

## Abstract

The molecular and intracellular signaling processes that control sleep and wake states remain largely unknown. A consistent observation is that the cyclic-AMP response element binding protein (CREB), an activity-dependent transcription factor, is differentially activated during sleep and wakefulness. CREB is phosphorylated by the cyclic AMP/protein kinase A (cAMP/PKA) signaling pathway as well as other kinases, and phosphorylated CREB (pCREB) promotes transcription of target genes. Genetic studies in flies and mice suggest that CREB signaling influences sleep/wake states by promoting and stabilizing wakefulness. However, it remains unclear where in the brain CREB is required to drive wakefulness. In rats, CREB phosphorylation increases in the cerebral cortex during wakefulness and decreases during sleep, but it is not known if this change is functionally relevant to the maintenance of wakefulness. Here, we used the cre/lox system to conditionally delete CREB in the forebrain and in the locus coereleus (LC), two regions known to be important for the production of arousal and wakefulness. We used polysomnography to measure sleep/wake levels and sleep architecture in conditional CREB mutant mice and control littermates. We found that forebrain-specific deletion of CREB decreased wakefulness and increased non-rapid eye movement (NREM) sleep. Mice lacking CREB in the forebrain were unable sustain normal periods of wakefulness. On the other hand, deletion of CREB from LC neurons did not change sleep/wake levels or sleep/wake architecture. Taken together, these results suggest that CREB is required in neurons within the forebrain but not in the LC to promote and stabilize wakefulness.

## Introduction

Mapping of the neural circuits that control wakefulness and sleep began over 60 years ago, with many studies delineating specific neural systems controlling sleep/wake cycles in the past two decades^1^. Electrophysiological and behavioral studies have led to an improved understanding of the neurotransmitter systems, pathways, and cell firing patterns that regulate these states. Wake promoting regions such as the locus coereleus (LC)^2^ send projections to the cerebral cortex and activity in these regions produces EEG desynchronization consistent with arousal and wakefulness^3^. This arousal producing system is balanced by sleep-promoting areas that induce sleep and suppress wakefulness^4^. These two mutually inhibiting systems act in concert with circadian and homeostatic processes to properly regulate sleep/wake states^5^. The combination of lesion studies and more recently, optogenetic and pharmacogenetic approaches has allowed the functional delineation of key sleep and wake regulatory circuits in the brain. However, much less is known about the intracellular processes underlying sleep/wake regulation^6^. Genetic manipulations and immunohistochemical measurements have identified the CREB signaling pathway and CREB-regulated transcription as potential regulators of sleep/wake states. The CREB signaling pathway has been extensively characterized in the context of synaptic plasticity and memory formation^7-13^. Interestingly, some studies also suggest that CREB signaling is also essential to the proper regulation of sleep/wake states. In flies, CREB activity is inversely related to rest^14^. Mice lacking two isoforms of CREB show reduced wakefulness and increased sleep^15^. However, it is unclear where in the brain CREB plays a critical role to drive wakefulness. CREB activity and expression of CREB downstream target genes are increased in the cortex following periods of wakefulness^16^.

These changes in the cortex require inputs from the LC^17^ and mice lacking dopamine beta hydroxylase (DBH), the enzyme required for norepinephrine synthesis, show reduced wakefulness and wake fragmentation ^18^. Here, we examine the role of CREB in the cortex and in the LC in promoting and stabilizing wakefulness using conditional knock out approaches and polysomnography.

## Methods

### Animals

Forebrain CREB cKO (FB CREB cKO) mice were produced by crossing animals expressing cre recombinase driven by the CaMKII promoter^19^ to mice carrying a floxed allele of Creb1^20^. Mice lacking CREB in noradrenergic neurons were produced by crossing mice expressing cre recombinase driven by the dopamine beta hydroxylase (DBH) promoter^21^ to mice carrying a floxed allele of Creb1AZ^20^. Experimental CREB conditional knock out animals were male and female mice homozygous for the floxed Creb1 allele and hemizygous for cre recombinase. Control mice were either hemizygous for cre recombinase or carried floxed alleles of Creb1. Animals were maintained on a 12 hour light/12 hour dark cycle with lights on (ZT 0) at 7:00 am. Food and water were available *ad libitum*. All animal care and experiments were approved by the Institutional Animal Care and Use Committee of the University of Pennsylvania and conducted in accordance with the National Institute of Health guidelines.

### Surgery

Animals were implanted with EEG and EMG electrodes under isoflurane anesthesia. Electrodes were held in place with dental cement (Ketac, 3M, St Paul, MN). Electrodes consisted of Teflon coated wires (Cooner wires, Chatsworth, CA) soldered to gold socket contacts (Plastics One, Roanoke, VA) and pushed into a 6-pin plastic plug (363 plug, Plastics One). The contacts were cemented to the plug using dental cement. Animals were connected to amplifiers using light-weight cables (363, Plastics One) attached to a rotating commutator (SLC6, Plastics One). All recordings were obtained using a parietal electrode (ML ±1.5mm, AP -4mm from bregma) referenced to an electrode over the cerebellum (1.5 mm posterior of lambda). Mice were allowed to recover from surgery for a minimum of 2 weeks. During the second week of recovery, mice were acclimated to the cables and to the recording chambers.

### EEG recordings and analysis

EEG/EMG signals were recorded over a 24 hour baseline period, during which animals were undisturbed. The next day, animals were sleep deprived (SD) for 6 hours, starting at lights on (ZT 0) using gentle handling^22^ and allowed to recover for 18 hours. EEG/EMG signals were sampled at 256 Hertz (Hz) and filtered at 0.5-30Hz and 1-100Hz, respectively with 12A5 amplifiers (Astro-Med, West Warwick, RI). Data acquisition and visual scoring was performed using SleepSign software (Kissei Comtec, INC, Japan). EEG/EMG recordings were scored in 4 second epochs as wake, NREM, or REM by a trained experimenter blind to genotype. Epochs containing movement artifacts were included in the state totals and architecture analysis, but excluded from subsequent spectral analysis. Spectral analysis was performed using a fast Fourier transform (FFT; 0.5-20Hz, 0.125Hz resolution). EEG spectra were computed over 24 hours. A few of the recordings had too many movement artifacts to process for FFT over 24 hours. These recordings were excluded from the 24-hour FFT analysis. NREM EEG spectra were computed in 2 hour windows for the baseline and 1 hour windows to the recovery period following SD. NREM slow wave activity (SWA), was normalized to the last 4 hours of the light phase of the baseline day for each animal as previously described^23^.

### Immunohistochemistry

Mice were anesthetized with isoflurane and transcardially perfused with ice cold PBS, followed by ice cold 4.0% paraformadelhyde using a peristaltic perfusion pump (Rainin Instruments, Oaklan, CA). Fixed brains were dissected, postfixed overnight and cryoprotected in 30% sucrose. Brains were frozen and cryosectioned coronally at a thickness of 30 um into PBS. Floating sections were blocked for 1 hour at room temperature in 2% normal donkey serum (NDS, Vector Laboratories, Burlingame, CA) in 0.3% Triton X-100 PBS. Sections were first processed for CREB-immunostaining with nickel ammonium sulfate intensification of DAB and then processed for tyrosine hydroxylase (TH). Sections were incubated overnight at 4C in 2% NDS, 0.1% Titon X-100 in PBS with anti-CREB primary antibody (1:3,000; 9197 Cell Signaling, Danvers, MA), 2 hours at room temperature with biotinylated donkey anti-Rabbit secondary antibody in 0.1% Triton X-100 PBS (1: 500; Jacskon Immunoresearch, Newmarket Suffolk, England) and 1.5 hours at room temperature with ABC in 0.1% Triton X-100 (1:500; Vector Laboratories, Burlingame, CA). Sections were washed 3 x 10 minutes with PBS between each incubation. The same procedure was followed for TH immunostains using anti-TH antibody (1:10,0000, Millipore, Billerica, MA). For fluorescent immunohistochemistry, FITC-conjugated anti-rabbit IgG secondary antibodie (1:1000, Chemicon, Temecula, CA) were used. Sections were mounted on slides and imaged using a microscope or confocal microscope. Cell counting and optical density measurements were conducted using ImageJ. 6 sections (3 from each side) from 3-4 animals were used for all quantifications of CREB levels.

### Quantitative real-time PCR

Cortical samples were collected into 500 uL of RNAlater (Ambion, Austin, TX) at ZT16 and flash frozen. All collections were done under red light to avoid any potential light pulse effects. The list of transcripts examined was based on previous reports showing that CREB is required for the expression of these genes^24^. Preparation of mRNA and cDNA synthesis were conducted as previously described^25^. PCR reactions were prepared in 96-well optical reaction plates (Applied Biosystems, Foster City, CA). Each well contained 10 uL of cDNA and 1uL of of Taqman probes and 9 uL of Taqman Fast Universal PCR Master Mix (Applied Biosystems, Foster City, CA). 6 biological and 3 technical replicates were used. Data were normalized to *Hprt1* (probe ID Mm0145399-m1, *Tuba4a* (Mm00849767-S1) and *Gapdh* (Mm99999915_g1). Fold change was calculated from the ΔCt cycle threshold values with corrections for standard curve data from each gene and housekeeping gene expression levels for each sample. For each sample, ΔCt was calculated against the mean for the sample set of that gene. Next, each of these values was corrected with the slope of the standard curve for the relevant probe to account for efficiency. This corrected value was normalized to the ΔCt from housekeeping genes for each sample to account for input variability. Fold change is equal to two raised to the difference between experimental and control Ct values. Student’s t-tests were used to compare groups.

### Statistics

Student’s t-tests were used to compare wake, NREM and REM sleep levels averaged over 24 hours. Multivariate analysis of variance (MANOVA) was used on the proportion of time spent in each state during the light phase, the dark phase and during the 8 x 3-hour time points across 24 hours, followed by Tukey studentized range tests to compare genotype groups. The same procedure was applied to the number of bouts and the average bout duration to compare groups. MANOVA was also used to compare the difference in total sleep time, NREM and REM sleep between the baseline and recovery periods following 6 hours of SD. Tukey studentized range tests were used to analyze raw EEG spectra for wake, NREM and REM sleep. Student’s t-tests were used to analyze theta peak frequency for EEG spectra. NREM SWA during the 4 x 2-hour bins was analyzed using repeated measures ANOVA followed by Tukey studentized range tests.

## Results

### CREB is required in excitatory neurons in the forebrain to drive and sustain wakefulness

Forebrain CREB conditional knock out mice (FB CREB cKO) were generated by crossing a cre-expressing line driven by the CaMKII promoter^19^ to animals carrying a floxed allele of Creb1^20^ (**Figure 1A**). Expression of *Creb1* and CREB target genes *Nr4a1* and *Gadd45b* was markedly reduced in the cortex of FB CREB cKO animals (Table 1). In contrast, expression of cAMP response element modulator (CREM), another member of the same family of transcription factors, was increased in conditional CREB mutants (Table1), which is consistent with previous manipulations decreasing *Creb1* expression^26^. CREB protein expression was substantially decreased in the cortex of FB CREB cKO (**Figure 1B-F**, p=0.005). We used EEG/EMG recordings to measure wake, non-rapid eye movement (NREM) and rapid eye movement (REM) sleep in conditional CREB mutant animals and control littermates. During the 24 h baseline period, animals were undisturbed and recording began at lights on (ZT0). FB CREB cKO animals showed decreased wakefulness over 24 hours (**Figure 2A**, t(26)=4.15, p=.0003) and during light and dark cycle (**Figure 2A**, F(2,25)=13.98, p<.0001).

**Table 1.**
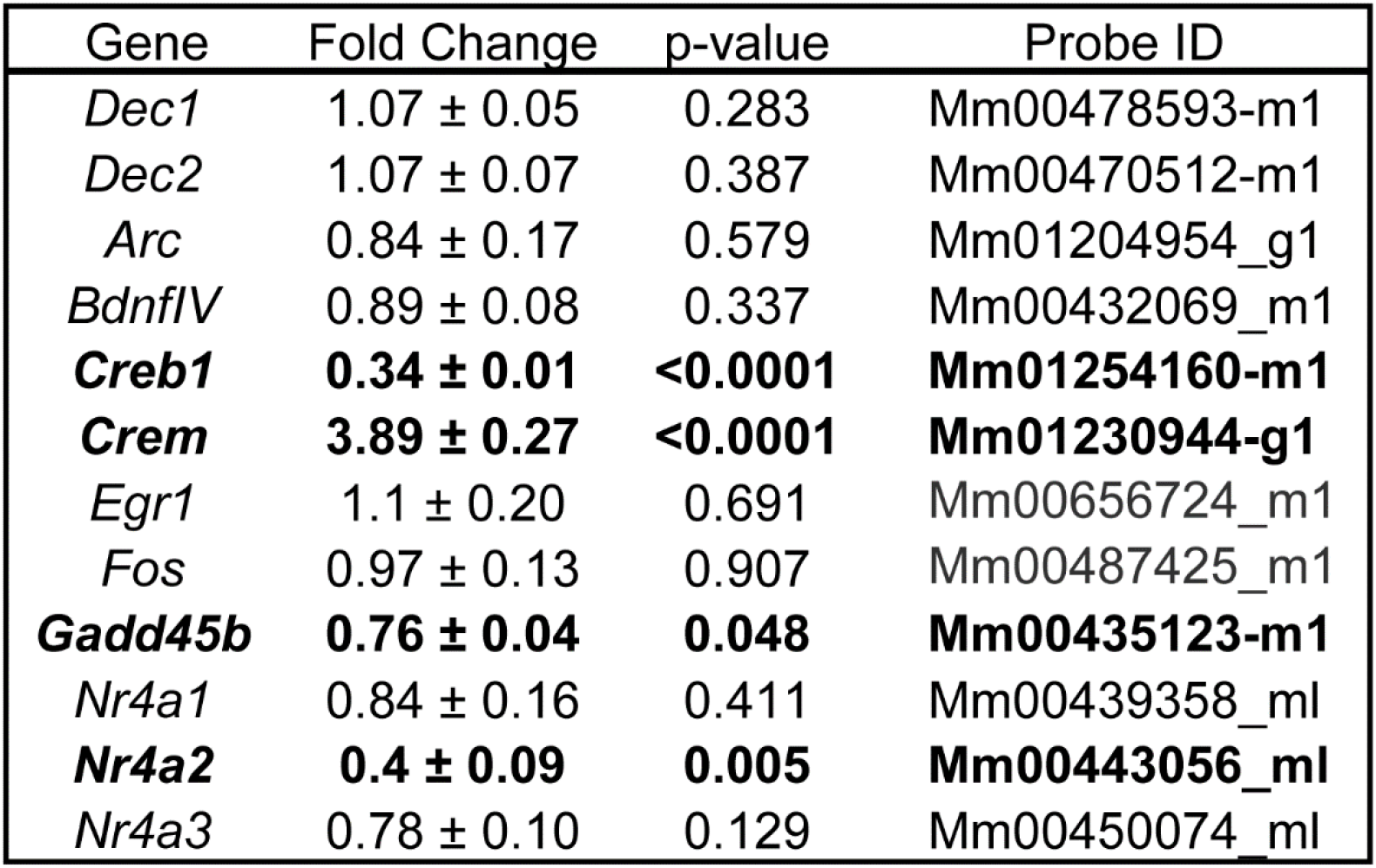
FB CREB cKO mice show reduced expression of CREB target genes in the cortex during the active phase. Tissue from cortex was collected at ZT 16 from FB CREB cKO and control mice (n=6). Gene expression was measured using qPCR. Transcripts changed by deletion of CREB are shown in bold. Mean ± s.e.m.

| Gene | Fold Change | p-value | Probe ID |
| --- | --- | --- | --- |
| *Dec1* | 1.07 ± 0.05 | 0.283 | Mm00478593-m1 |
| *Dec2* | 1.07 ± 0.07 | 0.387 | Mm00470512-m1 |
| *Arc* | 0.84 ± 0.17 | 0.579 | Mm01204954_g1 |
| *BdnfIV* | 0.89 ± 0.08 | 0.337 | Mm00432069_m1 |
| ***Creb1*** | **0.34 ± 0.01** | **<0.0001** | **Mm01254160-m1** |
| ***Crem*** | **3.89 ± 0.27** | **<0.0001** | **Mm01230944-g1** |
| *Egr1* | 1.1 ± 0.20 | 0.691 | Mm00656724_m1 |
| *Fos* | 0.97 ± 0.13 | 0.907 | Mm00487425_m1 |
| ***Gadd45b*** | **0.76 ± 0.04** | **0.048** | **Mm00435123-m1** |
| *Nr4a1* | 0.84 ± 0.16 | 0.411 | Mm00439358_ml |
| ***Nr4a2*** | **0.4 ± 0.09** | **0.005** | **Mm00443056_ml** |
| *Nr4a3* | 0.78 ± 0.10 | 0.129 | Mm00450074_ml |

**Figure 1.**
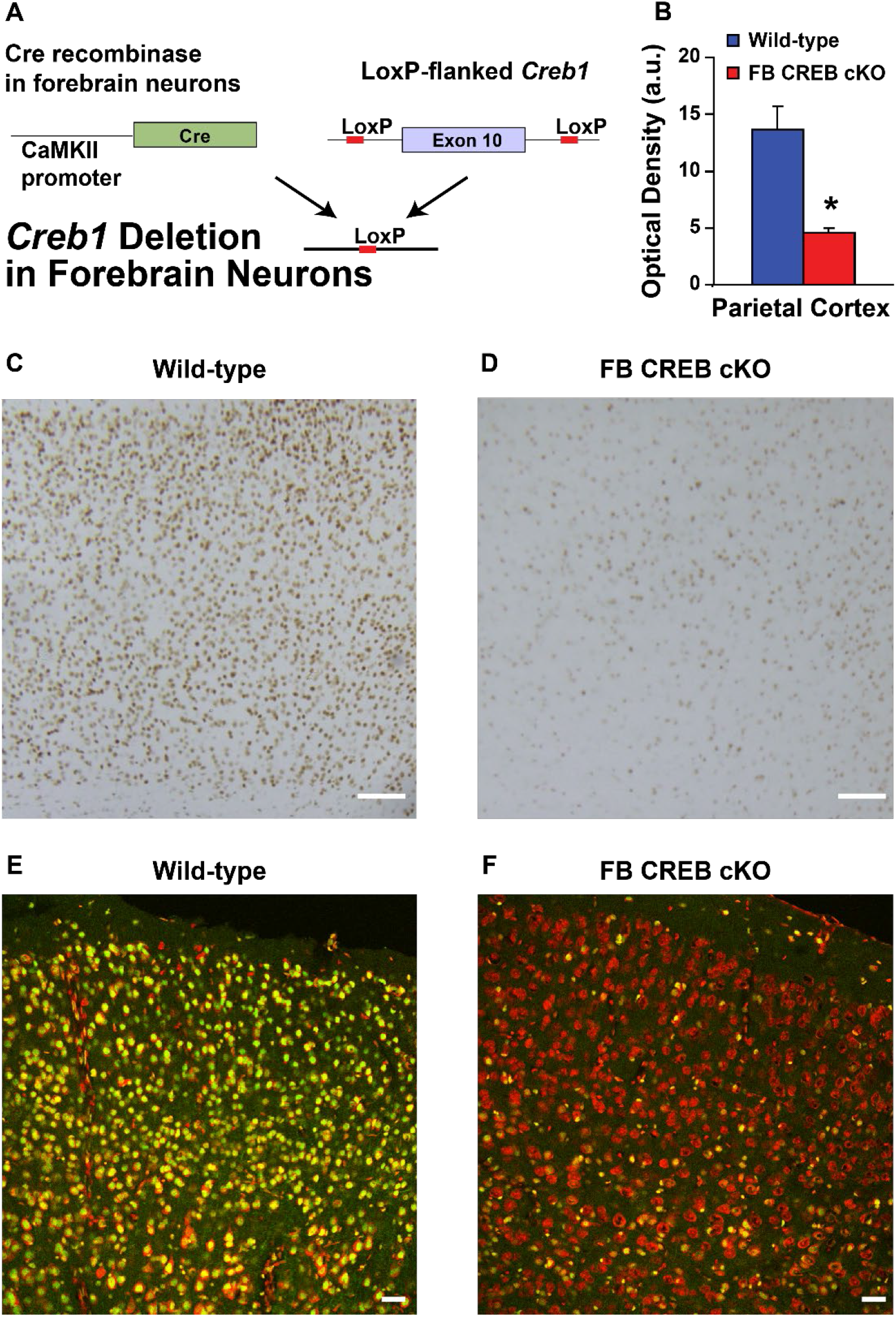
Forebrain CREB cKO animals show reduced CREB expression in cortex. **A**. Forebrain CREB cKO animals were generated using the cre/lox system. **B**. CREB protein levels were quantified using optical density in parietal cortex of FB CREB cKO and wild-type littermates. **C** and **D**. Representative immunohistochemical stains using anti-CREB antibody (brown nuclei) of cortex for wild-type (**C**) and FB CREB cKO (**D**) mice. **E** and **F**. Sections from wild-type (**E**) and FB CREB cKO (**F**) cortex tissue were imaged using fluorescent microscopy for CREB (green) and propridium iodide (red). * p<0.05.

**Figure 2.**
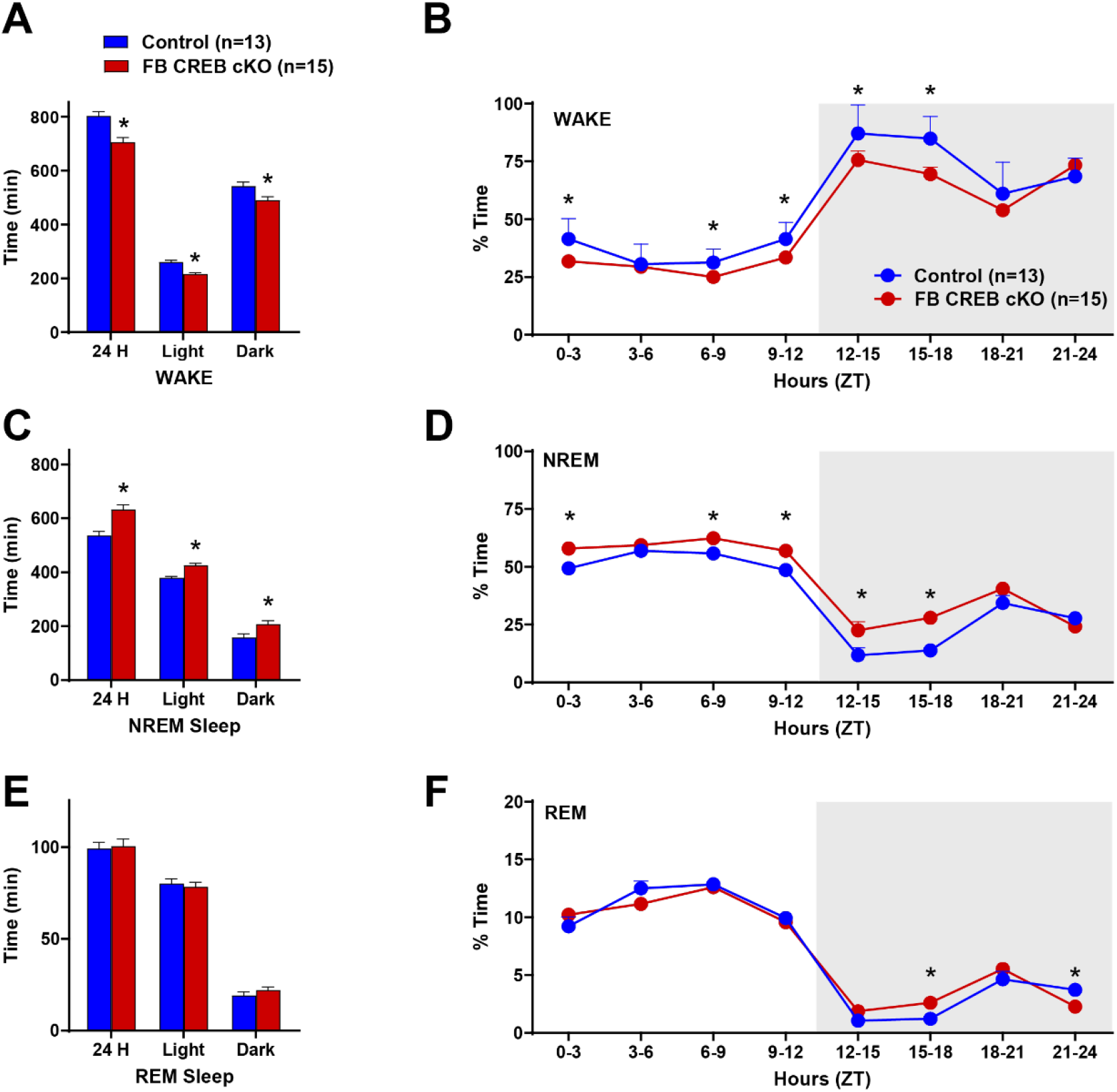
Deletion of CREB in forebrain neurons reduces wakefulness. **A**. Total time spent (min) in wakefulness over 24 hours and during the light and dark phases separately.**B**.Proportion of time in wake in 3 hour bins over a single 24 hour light/dark period. **C**. Total time in NREM sleep. **D**. Percentage of time in NREM sleep in FB CREB cKO and control mice. **E**. Total time spent in REM sleep over 24 hours and during each circadian period. **F**. Proportion of time in REM sleep displayed in 3-hour epochs across the light/dark cycle. Zeitgeber time (ZT) is represented on the x axis. Lights come on at ZT 0. * compares FB CREB cKO to controls (p< 0.05).

These changes in wakefulness were most pronounced during the early period of the dark phase (**Figure 2B**, F(8,19)=4.04,p=.0059; Tukey post-hoc tests). Decreased wakefulness was accompanied by increased NREM sleep in FB CREB cKO compared to control over 24 hours (t(26)=4.24, p=.0003) and during the light/dark phase (**Figure 2C**, F(2,25)=12.81,p=.0001). Analysis over 3-hour bins across light/dark revealed a large increase during the early part of the dark phase in particular (**Figure 2D**, F(8,19)=4.22, p=.0047. REM sleep levels were unchanged over 24 hours (t(26)=0.24, p=.8116) or during the light or dark phases (**Figure 2E**, F(2,25)=1.39, p=.2701). Multivariate analysis of variance (MANOVA) across 3-hour bins revealed more subtle changes in REM sleep during the dark phase (**Figure 2F**, F(8,19)=3.45, p=.0128).

### CREB is required in the forebrain to sustain wakefulness

Forebrain conditional knock out mice showed an increased number of wake bouts during the light and dark phase (**Figure 3A**, F(2,24)=5.38, p=.0017). In addition, the average duration of wake bouts was reduced in FB CREB cKO mice (**Figure 3B**, F(2,24=18.85, p<0.0001). The number of NREM bouts also increased in FB CREB cKO (**Figure 3C**, F(2,24)=8.55, p=.0016). However, the average duration of NREM bouts was unchanged by CREB deletion in the forebrain (**Figure 3D**, F(2,24)=2.17, p=.1364). The number of REM bouts was unchanged by CREB deletion (**Figure E**, F(2,24)=1.29, p=.2928) and the average duration of REM bouts were decreased in CREB conditional knock out mice only during the light phase (**Figure F**, F(2,24)=5.41, p=.0115, Tukey post-hoc test). Taken together, these results suggest that CREB is required in the forebrain to sustain wakefulness.

**Figure 3.**
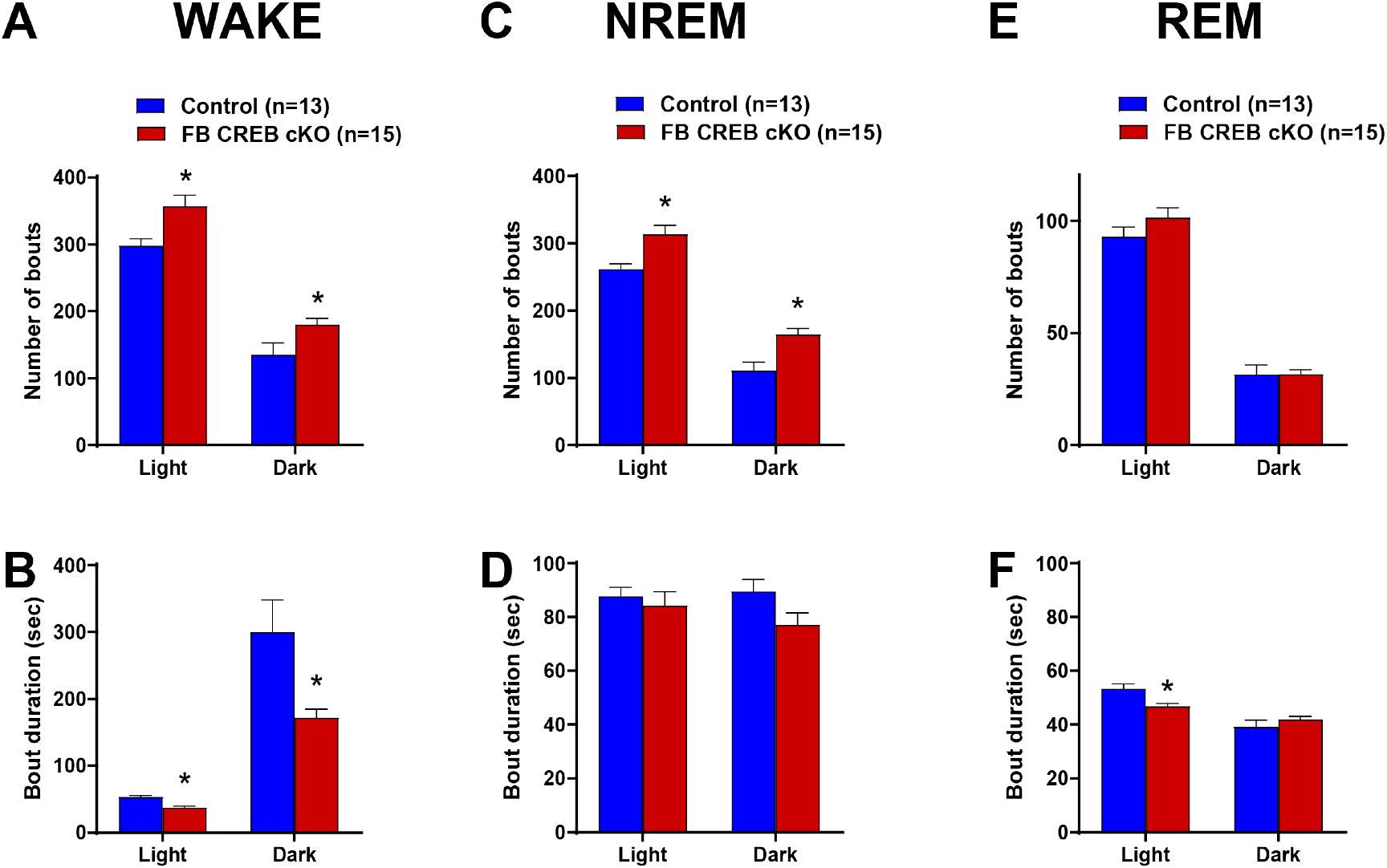
FB CREB cKO mice are unable to sustain wakefulness. Number of bouts and average bout duration for wake (**A** and **B**, respectively), NREM (**C** and **D**) and REM sleep (**E** and **F**) during the light and dark phase. Mean ± s.e.m.; * indicate p< 0.05.

### Deletion of CREB in the forebrain reduces sleep homeostasis

Following the baseline day, animals were sleep deprived for 6 hours using gentle handling^15,22,27^ starting at lights on and mice were allowed to recover for the subsequent 18 hours. FB CREB cKO animals showed lower NREM sleep rebound when released from 6 hours of sleep deprivation compared to wild-type littermates (**Figure 4A**, genotype: F(1,13)=6.02, p=.0290; time: F(7,13)=16.94, p<.0001; time by genotype: F(7,13)=2.81,p=.0907). In contrast, REM sleep rebound was not altered by deletion of CREB from forebrain neurons (**Figure 4B**, genotype: F(1,10)=0.05, p=.8296, time: F(7,13)=9.14,p<.0001; genotype by time interaction: F(7,13)=0.02, p=.9743). Slow wave activity (SWA), the power in the delta (0.5-4Hz) frequency range during NREM sleep, is a marker of sleep pressure and sleep intensity^23,28,29^. SWA was normalized to the last 4 hours of the light phase during the baseline period for each animal ^23,30,31^. During the baseline period, SWA was highest at lights on and decreased over the course of the light phase (**Figure 4C**, time: F(3,14)=31.04, p<.0001). FB CREB cKO showed reduced SWA during the baseline day (genotype: F(1,14)=16.33, p=.0012; time by genotype interaction: F(3,14)=2.62, p=.0635). Following sleep deprivation, SWA was highest when the animals were allowed to sleep (ZT 6) and decreased during recovery sleep **(Figure 4D**, time: F(3,14)=101.76, p<.0001). SWA was lower for FB CREB cKO (genotype: F(1,14=22.97,p=.0003); time by genotype interaction: F(3,14)=8.82, p=0.0001). These results indicate that CREB deletion reduces sleep rebound following sleep deprivation and prevents the accumulation of SWA, a marker of sleep pressure.

**Figure 4.**
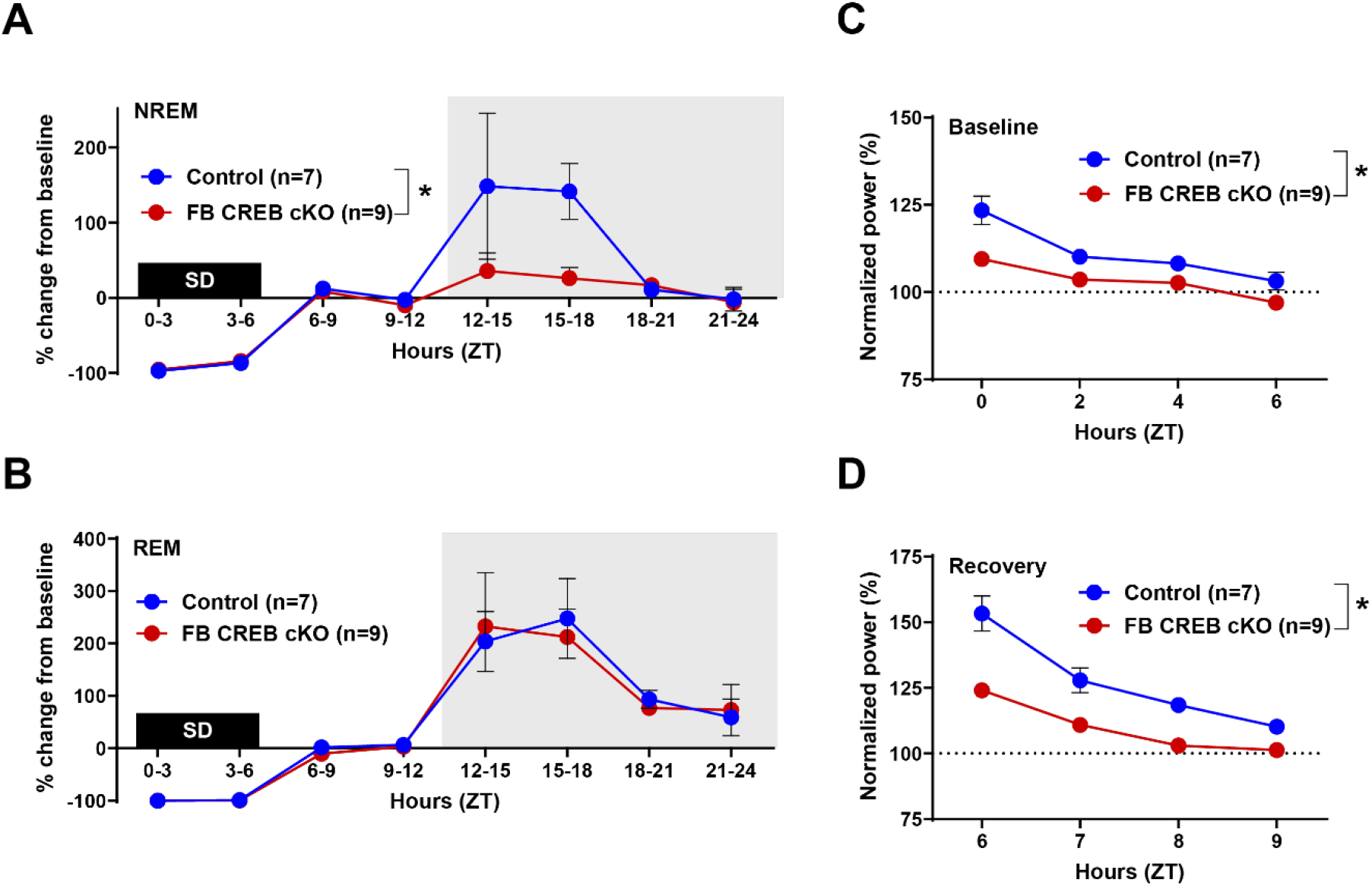
Forebrain deletion of CREB decreases sleep homeostasis. Animals were sleep deprived using gentle handling for 6 hours (ZT 0-6, labeled SD). Shown is percent change in NREM (**A**) and REM sleep time (**B**) between the recovery and baseline periods. **C**. Slow wave activity (SWA) expressed as percent baseline during baseline light period. **D**. SWA during recovery following 6 hours SD. Mean ± s.e.m., * p< 0.05.

### Deletion of CREB from forebrain neurons does not change EEG spectral content

We examined EEG spectral content using fast Fourier transform (FFT) of wake, NREM and REM sleep (**Figure 5**). In particular, we quantified power in the theta (6-10Hz) range during wakefulness, a marker of arousal, as well as power in the delta range during NREM sleep, an indicator of sleep pressure. We found no difference in absolute theta power during wake or REM sleep (**Table 2**). Delta power during NREM was also similar comparing conditional CREB knock out animals to controls.

**Table 2.**
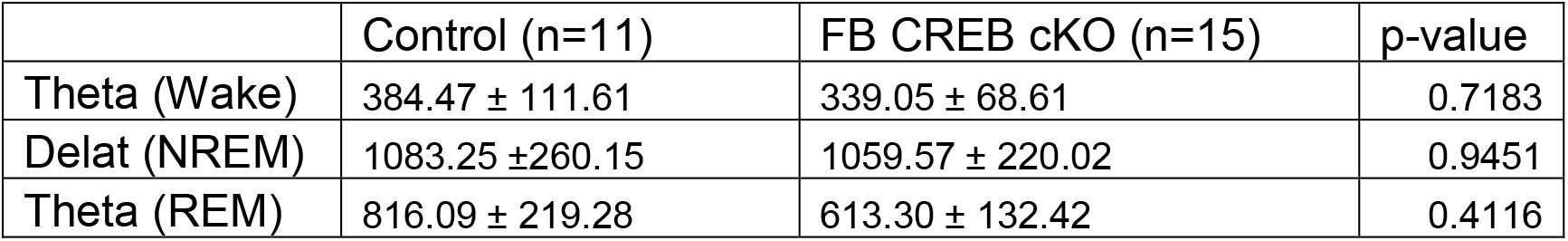
CREB deletion in the forebrain does not alter EEG sprectral power.

|  | Control (n=11) | FB CREB cKO (n=15) | p-value |
| --- | --- | --- | --- |
| Theta (Wake) | 384.47 ± 111.61 | 339.05 ± 68.61 | 0.7183 |
| Delta (NREM) | 1083.25 ± 260.15 | 1059.57 ± 220.02 | 0.9451 |
| Theta (REM) | 816.09 ± 219.28 | 613.30 ± 132.42 | 0.4116 |

**Figure 5.**
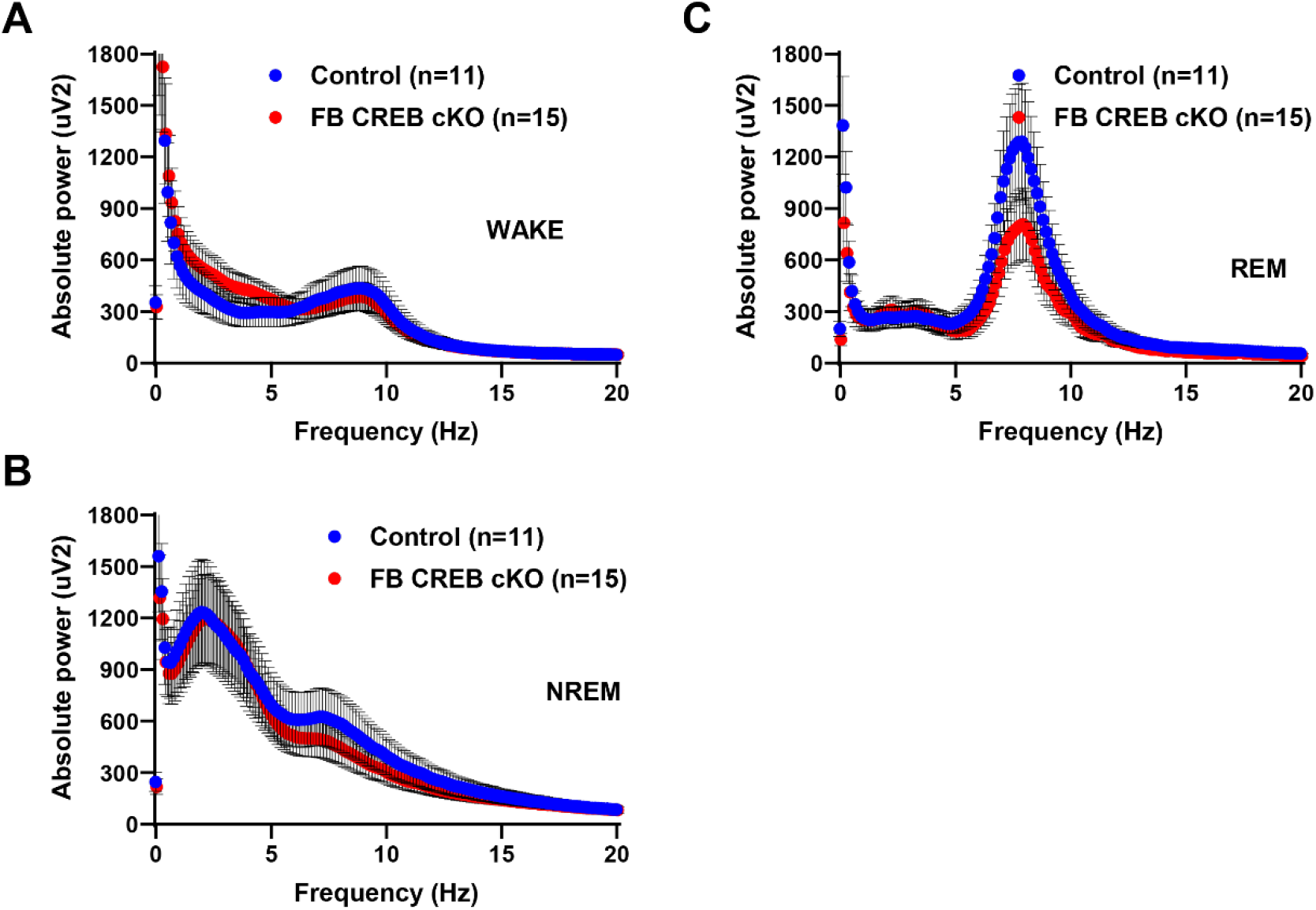
Deletion of CREB in forebrain does not alter EEG spectral content. EEG spectra were computed using FFT over 24 hours. Absolute power is shown for wake (**A**), NREM (**B**) and REM (**C**) sleep. Mean ± s.e.m.

A separate cohort of animals was used to examine circadian activity patterns in FB CREB cKO (n=6) and control (n=8) animals. Mice showed rhythmic activity patterns under 12h-12h light/dark conditions and under constant darkness. Deletion of CREB did not affect the period (control Tau=23.54±0.04; cKO Tau=23.64±0.08; p=0.202).

### Expression of Cre recombinase in forebrain neurons produces a subtle decrease in wakefulness

The control animals in the aforementioned studies did not express Cre recombinase. To address this potential caveat, we directly examined the potential impact of Cre recombinase expression alone on sleep/wake patterns. A separate cohort of animals was run through polysomnography to examine the potential impact of Cre recombinase expression on sleep/wake. Total wake time was reduced over 24 hours in animals expressing only Cre recombinase (**Figure 6A**, t(18)=.0345). However, total wake time over the dark and light phase was not changed by Cre recombinase expression (**Figure 6A**, F(2,17)=2.52, p=.1101). The proportion of time in wakefulness analyzed over 3 hour bins across the 24-hour light/dark cycle revealed no effect of Cre recombinase expression at any individual time window (**Figure 6B**, F(8,11)=0.97, p=.5070). Cre recombinase expression also increased NREM sleep overall (**Figure 6C**, t(18)=.0345), but when analyzed over the light and dark period, there was not significant difference between Cre-expressing and wild-type animals (**Figure 6C**, F(2,17)=2.53, p=.1090). The proportion of NREM sleep was also unaffected when examined in 3 hour bins (**Figure 6D**, F(8,11)=1.12, p=.4206). Total REM sleep time was unaffected by Cre recombinase expression (**Figure 6E**, 24 hours: t(18)=.9168; light/dark analysis: F(2,17)=0.60, p=.5576). The proportion of time spent in REM sleep was also intact comparing animals expressing Cre recombinase in the forebrain to controls (**Figure 6F**, F(8,11)=0.82, p=.6039).

**Figure 6.**
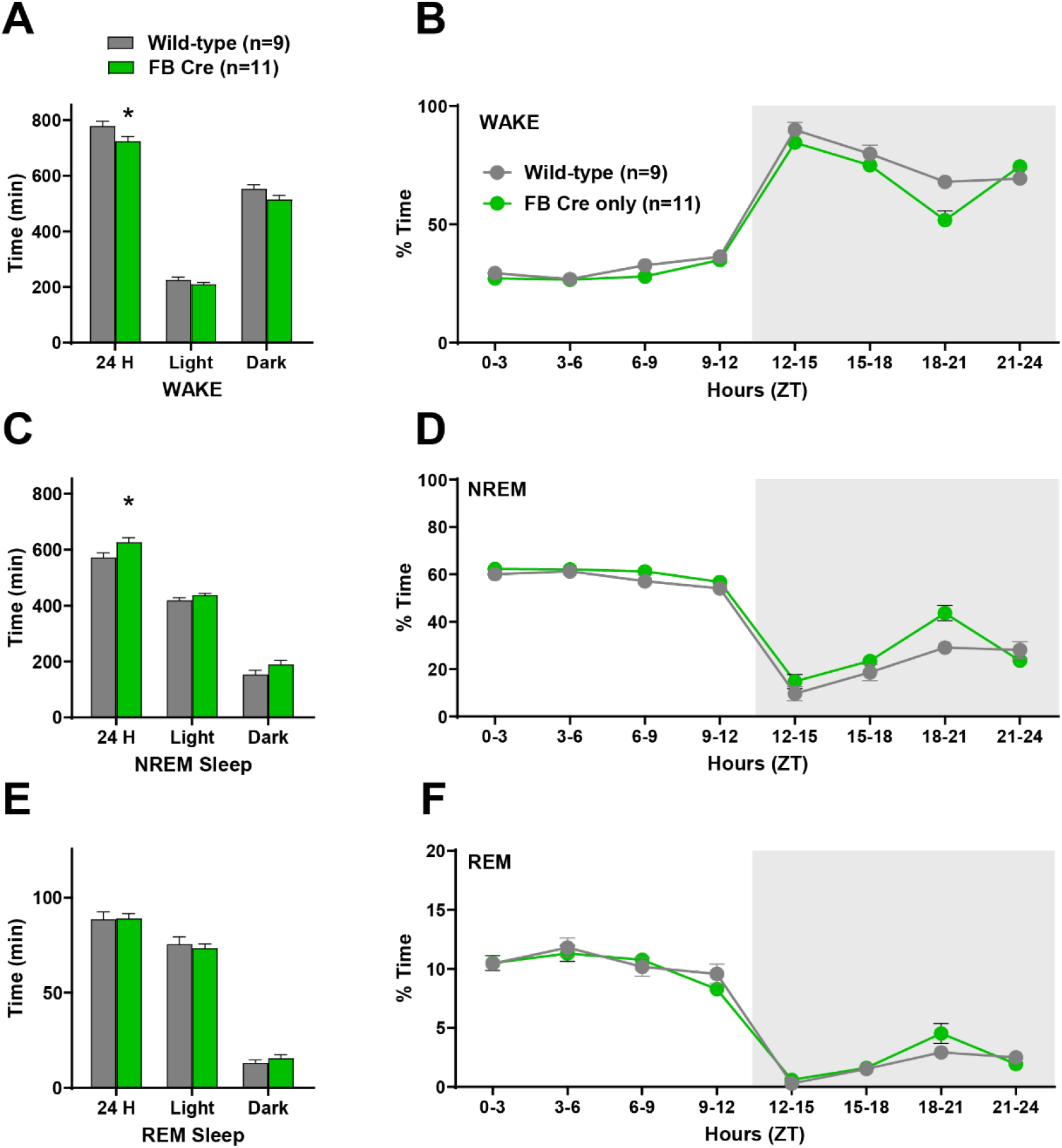
Expression of Cre recombinase in the forebrain reduces overall wakefulness. **A**. Total time spent (min) in wakefulness over 24 hours and during the light and dark phases separately. **B**. Proportion of time in wake in 3 hour bins over a single 24 hour light/dark period. **C**. Total time in NREM sleep **D**. Percent of time in NREM sleep **E**. Total time spent in REM sleep over 24 hours and during each circadian period. **F**. Proportion of time in REM sleep displayed in 3-hour epochs across the light/dark cycle. Zeitgeber time (ZT) is represented on the x axis. Lights come on at ZT 0. * indicate p<0.05.

### Expression of Cre recombinase in forebrain neurons does not impact sleep/wake architecture

The number of wake bouts was unaffected by Cre expression alone (**Figure 7A**, F(2,17)=1.30; p=.2982). The average duration of wake bouts was similar between animals expressing cre recombinase in the forebrain and wild-type controls (**Figure 7B**; F(2,17)=1.81, p=.1936). The number (**Figure 7C**; F(2,17)=0.93; p=.4128) and duration (**Figure 7D**; F(2,17)=0.18; p=.8374) of NREM bouts was unchanged by Cre recombinase expression in forebrain neurons. REM sleep architecture was also left intact by expression of Cre recombinase in the forebrain (**Figure 7E and 7F**; number of REM bouts: F(2,17)=2.14; p=.1484; REM bout duration: F(2,17)=0.47, p=.6340).

**Figure 7.**
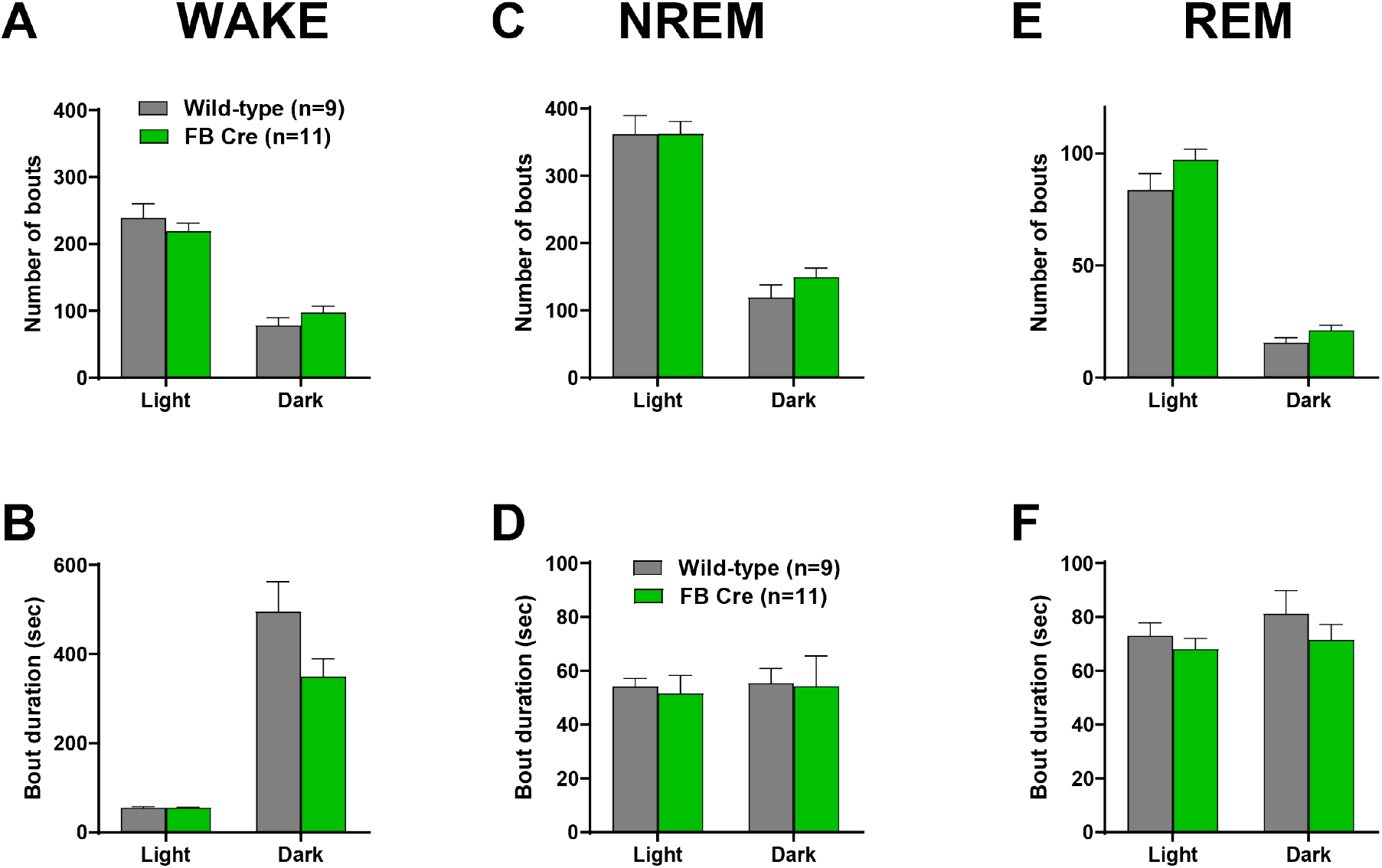
Expression of Cre recombinase in the forebrain does not alter sleep/wake architecture. Number of bouts and average bout duration for wake (**A** and **B**, respectively), NREM (**C** and **D**) and REM sleep (**E** and **F**) during the light and dark phase. Mean ± s.e.m.

### Expression of Cre recombinase does not impact EEG spectral content

We examined EEG spectral content using FFT of wake, NREM and REM sleep in animals expressing Cre recombinase in the forebrain compared to wild-type controls (**Figure 8**). In particular, we quantified power in the theta (6-10Hz) range during wakefulness, as well as power in the delta range during NREM sleep. There were no differences in absolute theta power during wake or REM sleep (**Table 3**). Delta power during NREM was also similar comparing conditional CREB knock out animals to controls (**Table 3**).

**Table 3.** EEG spectral power is not affected by expression of Cre recombinase in the forebrain

|  | Control (n=7) | FB Cre (n=9) | p-value |
| --- | --- | --- | --- |
| Theta (Wake) | 125.05 ± 21.79 | 120.02 ± 8.56 | 0.8174 |
| Delta (NREM) | 398.7 ± 78.16 | 428.65 ± 41.95 | 0.7249 |
| Theta (REM) | 215.59 ± 46.07 | 179.03 ± 13.98 | 0.4123 |

**Figure 8.**
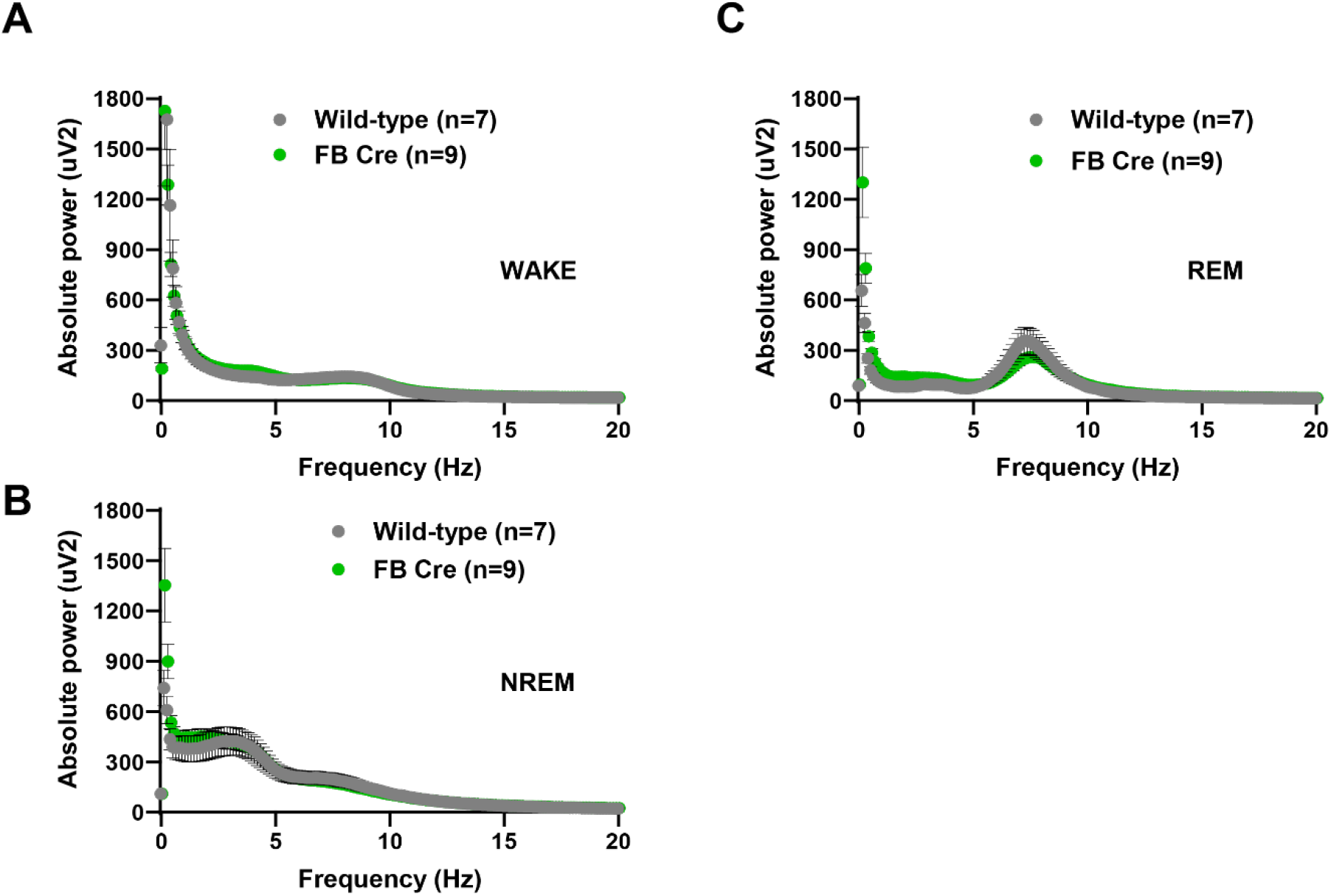
Expression of cre recombinase in the forebrain does not alter EEG spectral content. EEG spectra were computed using FFT over 24 hours. Absolute power is shown for wake (**A**), NREM (**B**) and REM (**C**) sleep. Mean ± s.e.m.

### CREB is not required in the locus coeruleus (LC) to sustain wakefulness

We used the DBH promoter^21^ to confer deletion of CREB to noradrenergic neurons (LC CREB cKO, **Figure 9A**). LC CREB cKO mice showed reduced expression of CREB in neurons that were positive for tyrosine hydroxylase (TH) in the LC (**Figure 9B-D;** 86% reduction, p=1.19 x 10^−5^). LC CREB cKO mice showed similar levels of wake compared to wild-type littermates over 24 hours (p=.4930) and during the light and dark phase (Figure **10A**, F(2,14)=0.25, p=.7803). NREM levels were also similar for both groups over 24 hours (p=.4397) and during the light and dark phase (**Figure 10B**, F(2,14)=0.31, p=.7383). REM sleep was also unchanged by deletion of CREB in LC neurons (**Figure 10C**, 24 hours, p=.9837, light/dark F(2,14)=0.01, p=.9950). LC CREB cKO showed similar levels of wake, NREM and REM sleep at each 3 hour time point across the light/dark cycle (wake: **Figure 10D**, F(8,8)=1.30, p=.3602, NREM: **Figure 10E**, F(8,8)=1.68, p=.2386, REM: **Figure 10F**, F(8,8)=0.43, p=.8702).

**Figure 9.**
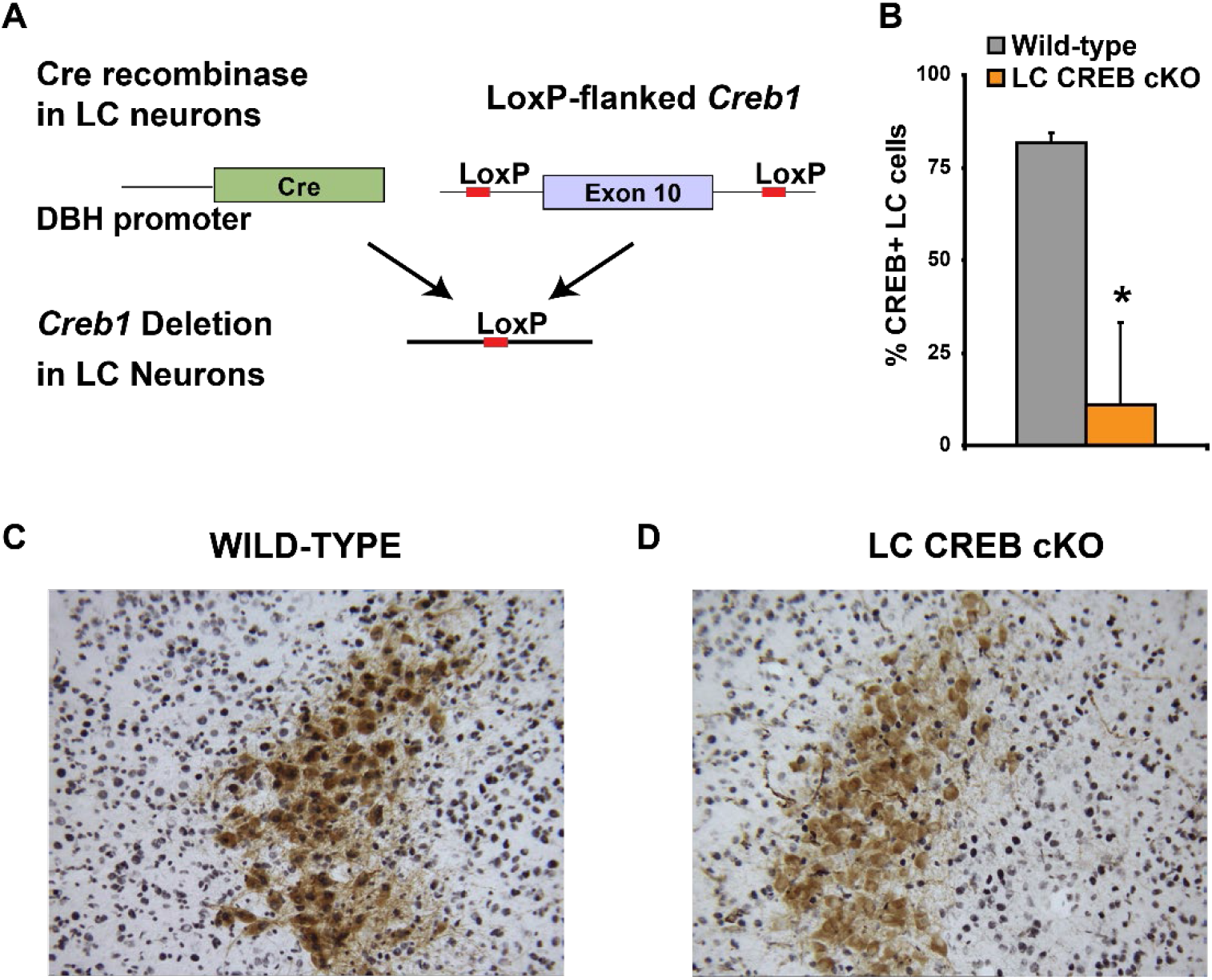
LC CREB cKO show reduced CREB expression in LC neurons. **A**. The cre/lox system was used to restrict deletion of CREB to noradrenergic neurons. **B**. Percentage of noradrenergic (TH positive) neurons also labeled for CREB is reduced in DBH CREB cKO animals. **C** and **D**. Representative photomicrograph of LC sections stained using an anti-TH antibody (brown cells) and anti-CREB antibody (blue nuclei) for wild-type (**C**) and mutant (**D**) mice. Mean ± s.e.m. * indicates p<0.05.

**Figure 10.**
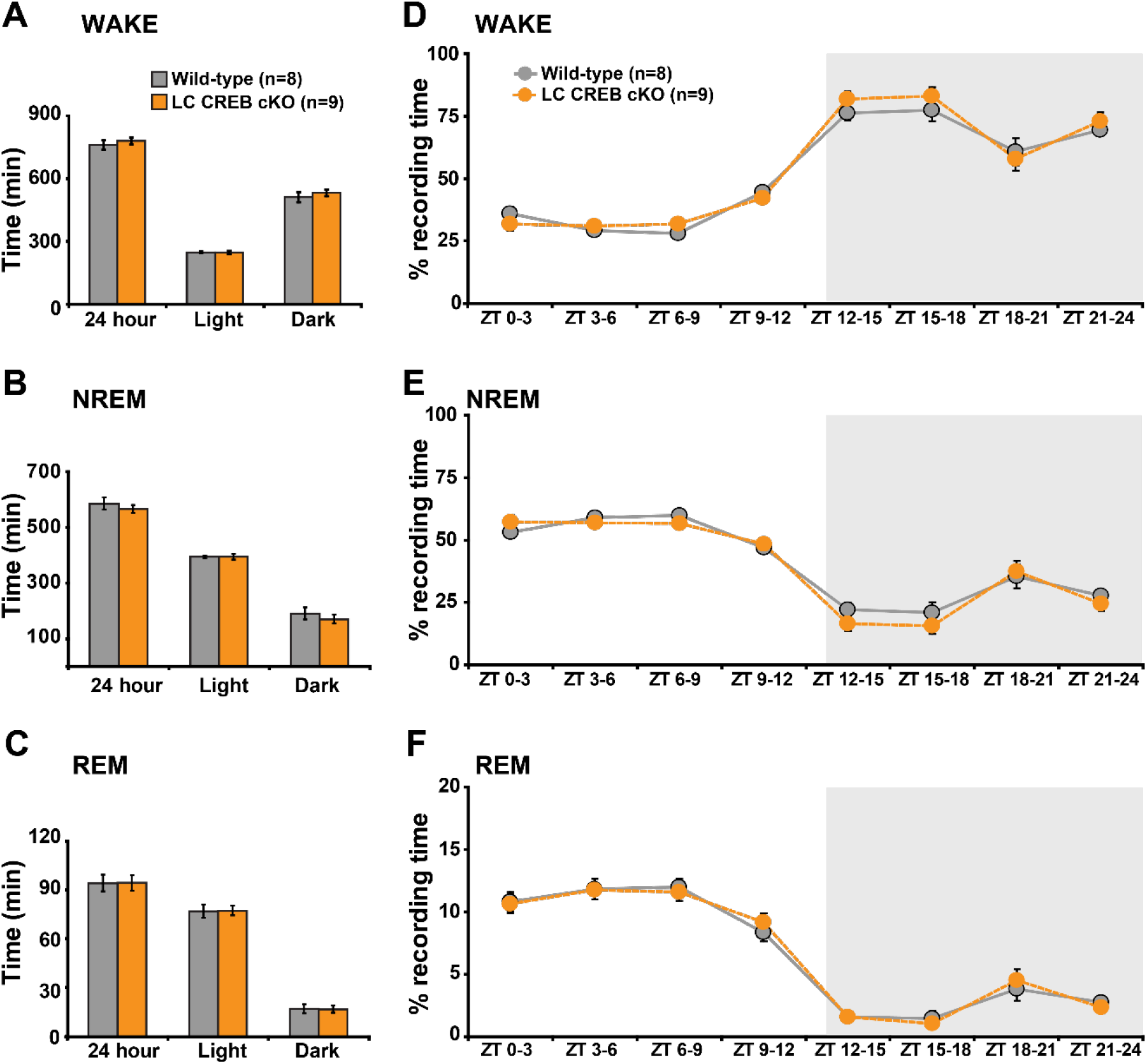
LC CREB cKO animals show normal levels of sleep/wake. Total time spent (min) in wake (**A**), NREM (**B**) or REM (**C**) sleep over 24 hours and during the light and dark phase. Percent time in wake (**D**), NREM (**E**) and REM sleep (**F**) across 24 h baseline period shown in 3 hour windows. Zeitgeber time (ZT) is represented on the x axis. Lights come on at ZT 0. Black line represents the dark period. Mean ± s.e.m.

Deletion of CREB from LC did not affect the number of bouts for wake (**Figure 11A**, F(2,14)=0.42, p=.6646) NREM (**Figure 11B;** F(2,14)=0.29, p=.7554) and REM sleep (**Figure 11C;** F(2,14)=0.42, p=.6630). The average duration for wake (**Figure 11C** F(2,14)=0.20, p=.8208), NREM (**Figure 11D;** F(2,14)=0.08, p=.9211) and REM sleep (**Figure 11E**, F(2,14)=1.23, p=.3255) was similar in LC CREB cKO compared to wild-type littermates. These results indicate that CREB is not required in the LC to promote wakefulness. We examined EEG spectral content using FFT of wake, NREM and REM sleep in conditional CREB knock out animals lacking expression of CREB in noradrenergic locus coeruleus neurons compared to wild-type controls (**Figure 12**). There were no difference in absolute theta power during wake or REM sleep (**Table 4**). Delta power during NREM was also similar comparing conditional CREB knock out animals to controls (**Table 4**).

**Table 4.**
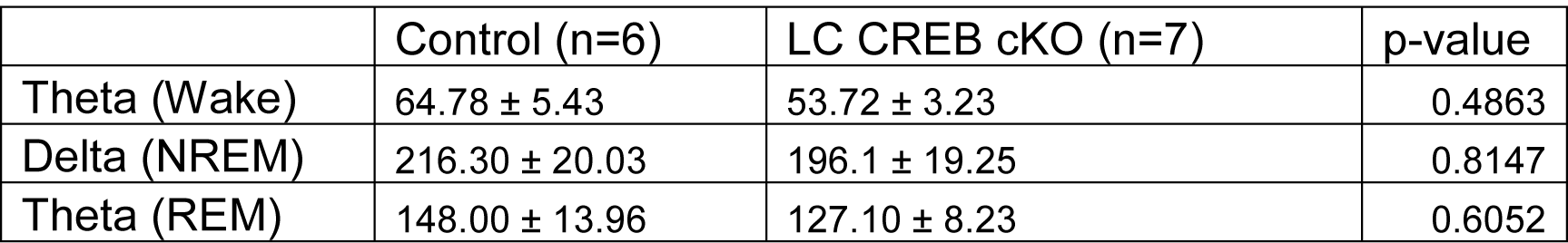
EEG spectral power is not affected by deletion of CREB from noradrenergic neurons

|  | Control (n=6) | LC CREB cKO (n=7) | p-value |
| --- | --- | --- | --- |
| Theta (Wake) | 64.78 ± 5.43 | 53.72 ± 3.23 | 0.4863 |
| Delta (NREM) | 216.30 ± 20.03 | 196.1 ± 19.25 | 0.8147 |
| Theta (REM) | 148.00 ± 13.96 | 127.10 ± 8.23 | 0.6052 |

**Figure 11.**
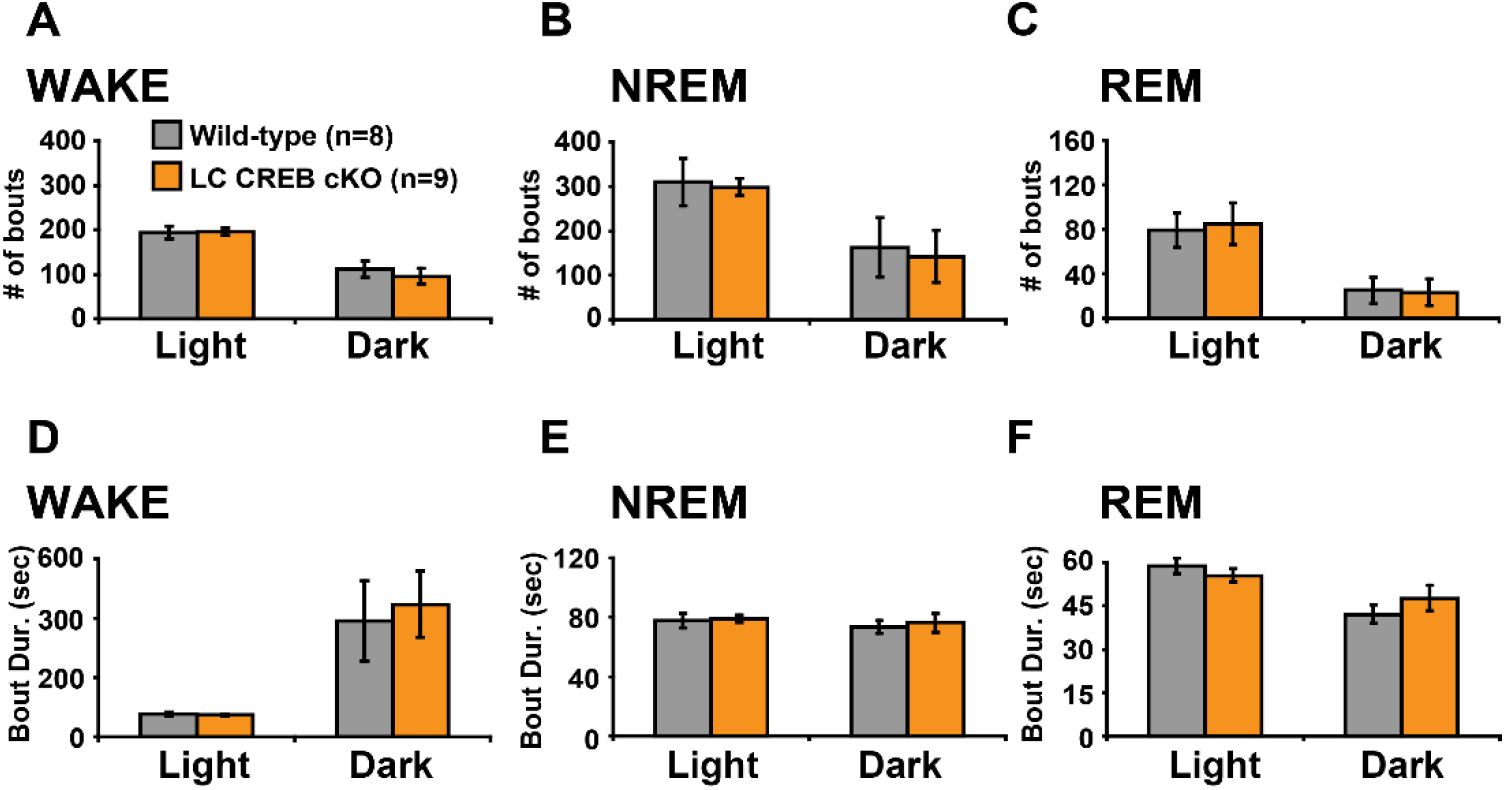
Deletion of CREB in LC does not disrupt sleep/wake architecture. Number of bouts for wake (**A**) NREM (**B**) and REM (**C)** and average bout duration for wake (**D**), NREM (**E**) and REM sleep (**F**). Mean ± s.e.m.

**Figure 12.**
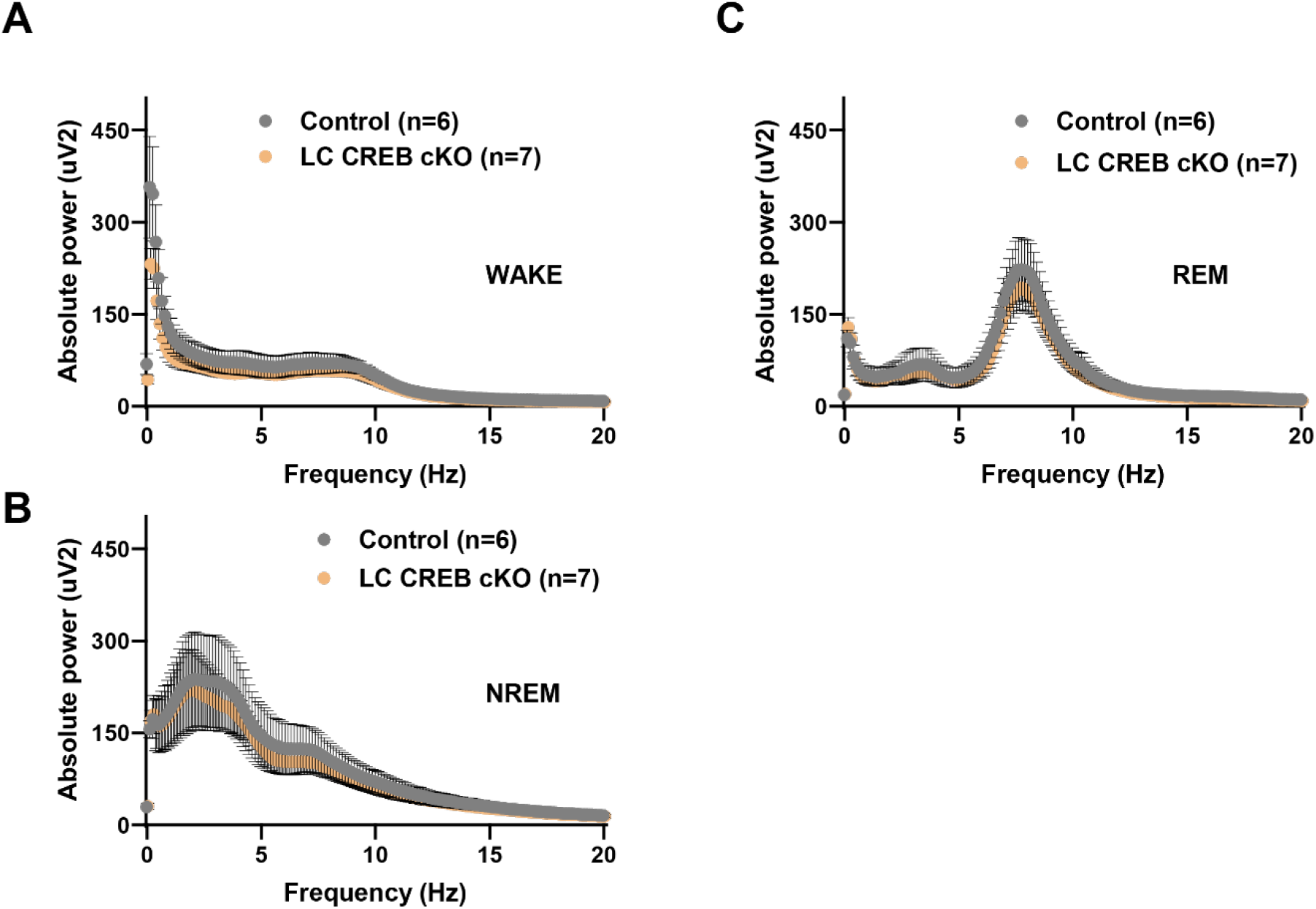
Deletion of CREB from noradrenergic neurons does not alter EEG spectral content. EEG spectra were computed using FFT over 24 hours. Absolute power is shown for wake (**A**), NREM (**B**) and REM (**C**) sleep. Mean ± s.e.m.

### Deletion of CREB from LC noradrenergic neurons does not alter sleep homeostasis

Sleep deprivation for 6 hours reduced NREM sleep and produced NREM sleep rebound during the recovery period (**Figure 13A**, effect of time: F(7,56)=39.15, p<.0001) similarly between CREB LC conditional knock out animals compared to controls (effect of genotype: F(1,8)=0.76, p=.4092; genotype by time interaction: F(7,56)=0.26, p=.8063). REM sleep was also increased similarly between LC CREB conditional knock outs and controls (**Figure 13B**, genotype: F(1,10)=0.43, p=.5282; time: F(7,70)=9.93, p<.0001; time by genotype interaction: F(7,70)=0.52, p=.7263). During the baseline period, SWA was highest at lights on and decreased over the course of the light phase (**Figure 13C**, time: F(3,42)=30.76, p<.0001) comparably between LC CREB cKO and controls (genotype: F(1,14)=0.57, p=.4642; time by genotype interaction: F(3,42)=0.56, p=.6438). Following sleep deprivation, SWA was highest when the animals were allowed to sleep (ZT 6) and decreased during recovery sleep **(Figure13D**, time: F(3,42)=35.63, p<.0001) with no impact of CREB deletion in LC neurons (genotype: F(1,14=1.00,p=.3335); time by genotype interaction: F(3,42)=1.32, p=.2809). These results indicate that CREB deletion in noradrenergic cells does not impact sleep rebound of SWA.

**Figure 13.**
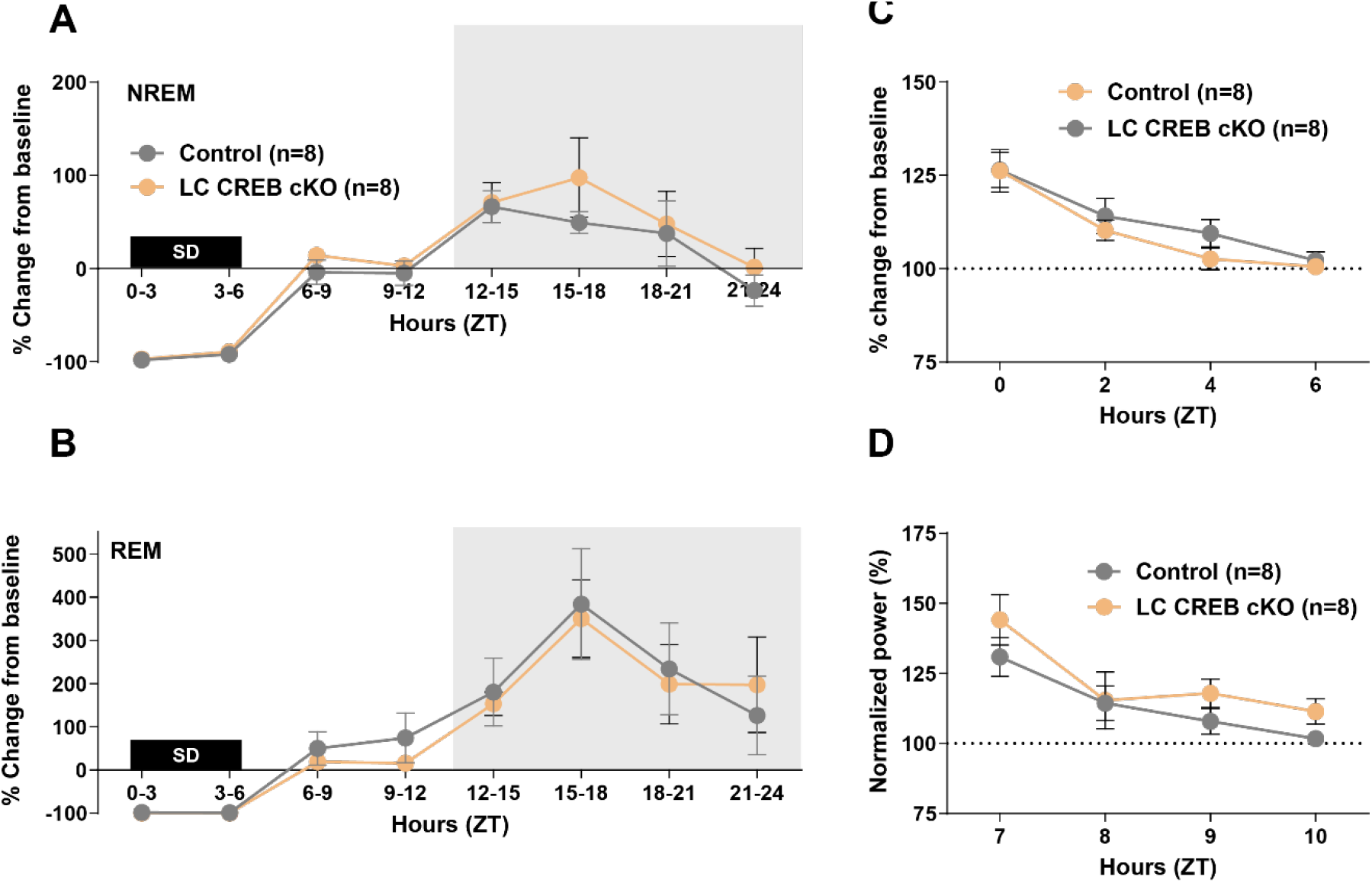
Deletion of CREB from LC neurons does not affect sleep homeostasis. Animals were sleep deprived using gentle handling for 6 hours (ZT 0-6). Shown is absolute change in NREM (**A**) and REM sleep time (**B**) between the recovery and baseline periods. Mean +/- s.e.m.

## Discussion

The goal of this study was to begin to functionally identify brain regions where CREB is important for the induction and maintenance of wakefulness. Activity in noradrenergic neurons of the locus coeruleus can desynchronize cortical EEG and produce arousal^2,3,32,33^. Wake promoting regions share the cerebral cortex as a common output and cortical activity is ultimately predictive of behavioral state^34,35^. We used conditional knock out approaches to separately test the role of CREB in these two key components of the wake promoting neural circuitry. Our results demonstrate that CREB is required in excitatory neurons in the forebrain but not in the LC to sustain wakefulness. The architecture of wakefulness was uniquely disrupted and fragmented by deletion of CREB in forebrain neurons. Interestingly, although the number of NREM bouts increased in forebrain CREB conditional knock out animals, NREM sleep architecture was largely unaffected suggesting that reductions in wakefulness were replaced by bouts of NREM sleep in these animals. We included a control experiment where Cre recombinase only was expressed in forebrain neurons. Cre-expressing animals showed a slight decrease in wakefulness and a concomitant increase in NREM sleep. These findings indicate that part of the phenotype associated with deletion of CREB in forebrain neurons could be due to Cre recombinase expression alone. However, it is noteworthy that animals expressing Cre recombinase in forebrain neurons had normal sleep and wake architecture, suggesting that the inability to sustain wakefulness in conditional forebrain CREB knock out animals can be solely attributed to CREB deletion in these neurons. The subtle but statistically significant change in sleep/wake levels in Cre recombinase-expressing animals underscores the need to thoroughly examine potential behavioral alterations elicited by expression of the Cre recombinase enzyme in the brain for each Cre-expressing mouse line. We decided not to undertake the same experiments for the LC conditional knock out animals because these mice showed no change in sleep/wake levels or architecture.

We used acute sleep deprivation to test the possibility that reduced and fragmented wakefulness resulted from increased sleep pressure in conditional CREB knock out animals. Chronic heightened sleep pressure is characterized by enhanced sleep homeostatic responses as shown in chronic sleep restriction experiments^36^ or in animal models with genetic manipulations resulting in increased sleep homeostasis^14,37,38^. Forebrain CREB conditional knock out mice had blunted NREM sleep rebound following 6 hours of sleep deprivation. Slow wave activity was reduced both during the baseline period and during recovery from acute sleep deprivation. It is possible that conditional CREB mutant mice did not reach the same levels of sleep pressure as control animals due to the increased opportunity to discharge slow wave activity during the active period. Interestingly, the slow wave activity levels reached by FB CREB cKO animals following 6 hours of enforced wakefulness are comparable to baseline SWA in control animals at lights on during the baseline day. These findings are consistent with the hypothesis that reduced SWA in CREB mutant mice stems from the wake fragmentation during the active period, which prevents the accumulation of SWA. Hence, our findings indicate that CREB is not directly involved in sleep homeostasis and that CREB does not serve a restorative function during sleep.

This phenotype of forebrain CREB conditional knock out animals is similar to that of animals constitutively lacking two isoforms of CREB^15^ and of animals lacking norepinephrine^18^. In the cortex, the increase in pCREB that occurs during wakefulness requires LC inputs^17^. Increased cortical norepinephrine during wakefulness^33^ could lead to increase pCREB via beta adrenergic Gs and cAMP/PKA activation and signaling. Another wake promoting mechanism that converges onto the cortex are peptidergic hypocretin/orexin inputs from the hypothalamus^35,39-42^. Activation of Gq-coupled orexin-2 receptors^43^could also lead to increased pCREB via intracellular calcium release. These neurons also send broad projections to other cortical layers and their activation in turn could result in broad cortical activity. Deletion of CREB in the cortex causes wake fragmentation, suggesting that the converging neural wake promoting systems require CREB downstream signaling to produce wakefulness.

Activity-dependent CREB transcription is regulated by a large set of signaling pathways. One example of this is protein kinase A, which has long been known to be a key part of CREB-mediated transcription involved in synaptic plasticity and memory formation^44,45^. In a drosophila model of Huntington’s Disease, animals showed increased wake and decreased sleep. This phenotype was at least partly dependent on increased PKA activity^46^. These findings are consistent with our current report and previous work in flies and mice supporting the idea that CREB is wake promoting. Another important regulator of CREB-mediated transcription is the protein kinase ERK (extracellular signal-regulated kinase). Deletion of *Erk1* and *Erk2* genes that encode ERK protein lead to increased wake and decreased sleep. ERK phosphorylation is thought to be critical for transducing waking experience and associated plasticity into sleep. Thus the disruptions in sleep and wake caused by ERK manipulations have been interpreted as deficits in wake-induced plasticity which ultimately impacts the quantity and quality of sleep ^47,48^. Taken together, these studies demonstrate that CREB signaling as well as several upstream regulators of CREB activity are involved in the regulation of sleep and wakefulness. Further studies are needed to establish the region specificity of these observations and the complex interactions of these molecules in the context of sleep/wake and related neural plasticity.

Consistent with previous reports, we found that deletion of CREB in the forebrain was accompanied by compensatory increased expression of cAMP response element modulator (CREM) ^20,21,26,49-51^. CREM is nearly undetectable in the brain under normal conditions and this marked compensatory increase has been shown to partly restore CREB function^20^. We also found reduced expression of CREB target genes *Nr4a2* and *Gadd45b* in CREB conditional knock out mice. Interestingly, activity-dependent expression of these transcripts requires CREB activity in the hippocampus and striatum^24^. Further studies are needed to determine whether these changes in gene expression are functionally relevant to the maintenance of wakefulness. This work suggests that the arousal system efferents to the forebrain promote wakefulness via activation of CREB signaling. Overall, our combined results have emphasized the importance of CREB signaling within forebrain neurons in promoting and sustaining wakefulness. Further studies are needed to refine where in the forebrain CREB is required for the maintenance of wakefulness and which effector genes mediate arousal in these cells.

## Acknowledgements

We dedicate this article to Dr. Günther Schütz, who was a pioneer in neuroscience and molecular biology research.

